# Assessing positive selection in centromere-associated kinetochore proteins across Metazoan groups

**DOI:** 10.64898/2026.02.13.705784

**Authors:** Hope M. Healey, L. Enrique Gomez, Sofia I. Sheikh, Benjamin R. Camel, Andrew A. Forbes, Kirstin N. Sterner, Emily A. Beck

**Affiliations:** Institute of Ecology and Evolution, University of Oregon, Eugene, OR, USA; Department of Anthropology, University of Oregon, Eugene, OR, USA; Department of Anthropology, Washington University in St. Louis, St. Louis, MO, USA; Department of Biology, University of Iowa, Iowa City, IA, USA; Department of Ecology and Evolution, University of Chicago, Chicago, IL, USA; Department of Molecular Biosciences, University of Kansas, Lawrence, KS, USA; Data Science Department, University of Oregon, Eugene, OR, USA

**Keywords:** positive selection, adaptive evolution, centromere drive, kinetochore, condensin

## Abstract

Centromeres are comprised of long stretches of repetitive DNA that evolve rapidly in organisms across the tree of life. Consistent selfish centromere evolution can also have cascading effects − driving rapid evolution in interacting kinetochore proteins – possibly to maintain centromere-kinetochore compatibility. Effects of selfishly evolving centromeres on interacting proteins are most heavily studied in the inner kinetochore and assembly proteins including the constitutive centromere-associated network proteins CENP-A and CENP-C with some exploration of the extended effects to other kinetochore-associated protein complexes. While rapid evolution of the centromere has been broadly studied in many organisms, studies assessing positive selection in centromere-associated kinetochore proteins have largely focused on *Drosophila*. Here, we tested the hypothesis that signatures of positive selection would be present in outer kinetochore and condensin genes in diverse animal groups. We selected two protein complexes −the Condensin I complex and the Mis12 Complex – to test for positive selection in parasitic wasps, two groups of ray-finned fishes (including the amazon molly an asexual diploid exempt from centromere drive), and two groups of primates. We did not find selection using any test in any protein in the amazon molly but did find sporadic positive selection in proteins in both complexes across all groups.

## Introduction

Centromeres are essential chromosomal regions largely comprised of tandem satellite repeats that play key roles in spindle attachment, chromosome segregation, and cell division in both mitosis and meiosis (Vig 1982; Bernard et al. 2001; Nasmyth 2002). While centromeres are highly conserved in function, they are highly variable in size, structure, and localization across taxa (Balzano and Giunta 2020; Talbert and Henikoff 2020). Centromeres also consistently evolve rapidly, and often selfishly, despite their need for functional constraints – a discordance termed “the Centromere Paradox” (Henikoff et al. 2001; Malik and Henikoff 2001; Bayes and Malik 2008; Brown and O’Neill 2014). One plausible explanation for The Centromere Paradox is centromere drive, a type of meiotic drive where larger centromeres are preferentially selected for incorporation into heritable oocytes during the first asymmetric division of female meiosis. Preferential incorporation of large centromeres into the oocyte results in transmission bias towards larger centromeres (Malik, 2009; Kursel & Malik, 2018; Dudka and Lampson 2022). This transmission bias drives selfish evolution of increasingly larger centromeres until a tipping point is reached; when centromeres become too large, leading to detrimental effects like formation of ectopic centromeres or improper spindle attachment subsequently constraining centromere size evolution (Heun et al. 2006).

Centromere drive requires more than just the rapid evolution of centromeric sequences. For centromeres to physically increase in size they also must recruit more inner kinetochore proteins (Heun et al. 2006). Increasing centromere size therefore requires recognition of the centromere by inner kinetochore proteins. Thus, as centromeres evolve, interacting components must also rapidly evolve in a Red Queen-like scenario (Malik and Henikoff 2001; Malik and Henikoff, S. 2002; Presgraves and Stephan 2007; Mensch et al. 2013). These coevolutionary dynamics can be identified genetically via signatures of positive selection. Signatures of positive selection are most commonly reported in the genes encoding inner kinetochore proteins CENP-A (Cid in Drosophila, CenH3 in plants, and Cse4 in yeast) and CENP-C, which are essential for kinetochore assembly, chromosome segregation, and passing mitotic checkpoints (Kwon et al. 2007; De Rop et al. 2012).

While *Drosophila* and yellow monkeyflowers (*Mimulus guttatus*/*M. nasutus*) have been primary models for centromere drive (Dudka and Lampson 2022; Kyriacou and Heun 2023), positive selection in genes encoding CENP-A and/or CENP-C has been identified across many plant species (Cooper and Henikoff 2004; Talbert et al. 2004; Finseth et al. 2015; Finseth et al. 2021), mice (Kumon et al. 2021; Arora and Dumont 2024), non-primate mammals (Pontremoli et al. 2021) and primates (Schueler et al. 2010; Arora and Dumont 2024). In ray-finned fishes, *cenp-a* has also exhibited signatures of positive selection, (Fountain and Kral 2011; Abbey and Kral 2015) but *cenp-c* has either been lost, or is evolving so rapidly that it is not readily identifiable in existing genome annotations (Kral 2016). In *Drosophila*, widespread signatures of positive selection have also been identified in genes encoding proteins interacting with the inner kinetochore. Specifically, signatures of positive selection have been reported in kinetochore protein chaperone genes like *cal1* (Rosin and Mellone 2016; Rosin and Mellone 2017) and in genes encoding components of larger interacting protein complexes i.e., Condensin I and Condensin II (Beck and Llopart 2015; King et al. 2019). In other species, including various groups of eutherian mammals, positive selection was found in genes encoding components of the Mis12 Complex (Pontremoli et al. 2021) and other kinetochore-associated proteins (Arora and Dumont 2024).

We hypothesized that positive selection in the centromere and inner kinetochore, would be accompanied by positive selection in outer kinetochore and condensin genes across animal phyla. To test this hypothesis, we sampled three groups of animals: insects, fish, and primates. Specifically, we selected a second group of insects outside *Drosophila* (parasitic wasps in family Braconidae), as well as two groups of ray-finned fishes (families Cichlidae and Poeciliidae), and two groups of primates from the parvorder Catarrhini (families Hominidae and Cercopithecidae). Within Poeciliidae, we included the amazon molly (*P. formosa)* – an asexual diploid that reproduces via gynogenesis (Hubbs and Hubbs 1932; Schlupp 2005; Schlupp et al. 2007) and should not experience centromere drive or subsequent cascades of positive selection to kinetochore proteins.

To expand assessment of selection outside the centromere, we assessed inner kinetochore genes *cenp-*a and *cenp-c*, and genes encoding components of two protein complexes associated with the inner kinetochore to test for positive selection. The Condensin I complex is physically associated with CENP-A and includes 5 proteins (SMC2, SMC4, CAP-G, CAP-D2, CAP-H). The Mis12 Complex is physically associated with CENP-C and contains 4 proteins (MIS12, DSN1, PMF1, and NSL1) (Figure 1). We did not anticipate finding positive selection in every gene as centromeres are known to drive positive selection in interacting proteins differently across species (Kyriacou and Heun 2023). However, we did anticipate finding positive selection in all groups. We found positive selection in a subset of these proteins in each group but did not find pervasive patterns consistent with those reported in *Drosophila*.

**Figure 1.**
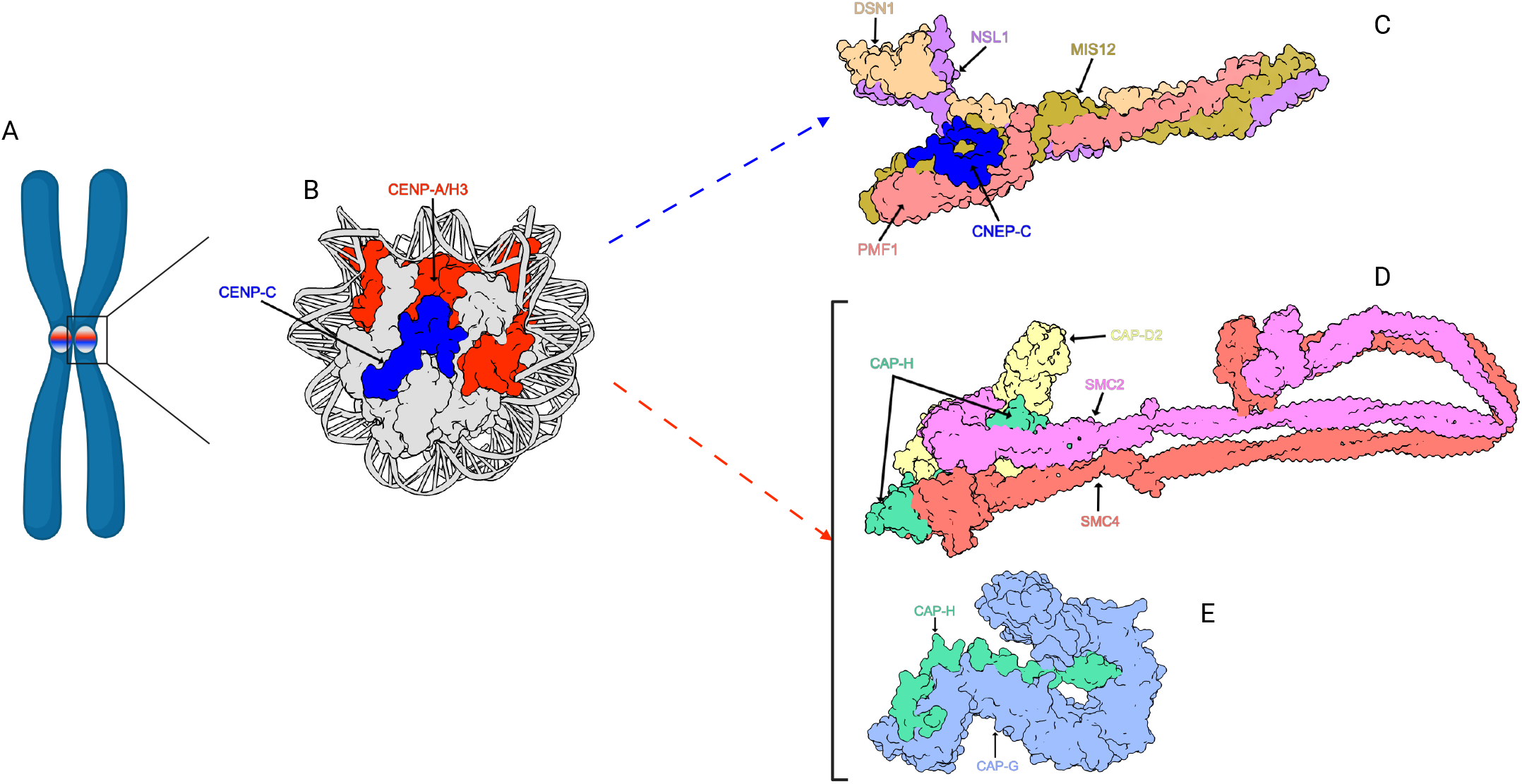
Protein complex schematic and graphical hypothesis of cascading selection. (A) Representation of chromosome highlighting the centromeric region. (B) CENP-A, CENP-C, Nucleosome complex at the centromere. (C) MIS12/CENP-C Complex. (D) SMC2/SMC4/NCAP-D2/NCAP-H Partial structure of Condensin I interacting with CENP-A. (E) NCAP-G and NCAP-H Partial Structure of Condensin I interacting with CENP-A. Subunits are colored differently and labeled by name.

## Materials and methods

### Sequences used

To obtain individual gene sequences from selected species in representative groups (insects, ray-finned fishes, and primates) we used sequencing data from various sources. For insects, we included a single group of wasps from the family Braconidae to be compared to published analyses of the *Drosophila melanogaster* subgroup (Beck and Llopart 2015). We collected Braconidae sequences from *Cotesia vestalis, Fopius arisanus*, and *Microplitis demolitor, Diachasma alloeum* from unpublished genomes and additional sequences of *Diachasma alloeum, D. ferrugineum, D. muliebre, Diachasmimorpha mellea* from genomes generated by the Forbes Lab (Tvedte et al. 2019; Tvedte et al. 2020). All sequences from unpublished genomes will be available on NCBI at the time of publication. For loci without public accessions (typically those derived from unpublished or incompletely annotated assemblies), orthologs were identified using a BLAST-based approach. Well-annotated Braconidae genes— verified against *Drosophila* orthologs from FlyBase—were used as nucleotide and protein queries to search target assemblies, and putative hits were evaluated for sequence homology and exon–intron structure and verified with NCBI ORFfinder. Searches were conducted either with NCBI BLAST (for publicly annotated genomes) or with Geneious v9.1.8 BLAST searches against custom databases constructed from genome assemblies using default megablast parameters. For ray-finned fishes, we included two groups: one from the family Cichlidae (*Amphilophus citrinellus, Astatotilapia burtoni, A. calliptera, Maylandia zebra, Neolamprologus brichardi, Oreochromis aureus*, and *Pundamilia nyererei*) and a second from the family Poeciliidae (*Gambusia affinis, Poecilia Formosa, P. latipinna, P. Mexicana, P. reticulata, Xiphophorus couchianus, X. maculatus*). We obtained all sequences from Ensembl version 98 (www.ensembl.org) (1 to 1 orthologs with high confidence; accessed 12/2019-02/2020) (Accession IDs available in Table S1). For primates we used two groups: one from the family Hominidae (*Gorilla gorilla gorilla, Homo sapiens, Nomascus leucogenys, Pan paniscus, P. troglodytes, Pongo abelii*) and the second from the family Cercopithecidae (*Piliocolobus tephrosceles, Colobus angolensis, Rhinopithecus sp*., *Chlorocebus sabaeus, Cerocebus atys, Mandrillus leucophaeus, Theropithecus gelada, Papio anubis, Macaca fascicularis*.). For the family Cercopithecidae, we had an abundance of sequence data and selected individual species as representatives for specific groups. Specifically, for *Rhinopithecus* we selected either *R. roxellana* or *R. bieti*. Lastly, we included either *Piliocolobus* or *Colobus* but not both. We obtained all primate sequences from Ensembl version 98 (1 to 1 orthologs with high confidence; accessed August 2022) or NCBI when not available or problematic on Ensembl. (Accession IDs available in Table S1).

Like others, we were unable to identify *cenp-c* in any of our fish (Kral 2016). We were also unable to find *dsn1* in fish and *smc4* in wasps. We also removed *cenp-*a from analysis in wasps as orthology was ambiguous and we did not want to inflate sequence divergence estimates by comparing across incorrect orthologs. In some specific species we were also missing individual protein annotations which could not be included in our analysis (Figure S1). For ease of reading, we use the human nomenclature for each gene/protein throughout the manuscript.

**Supplementary Figure 1.**
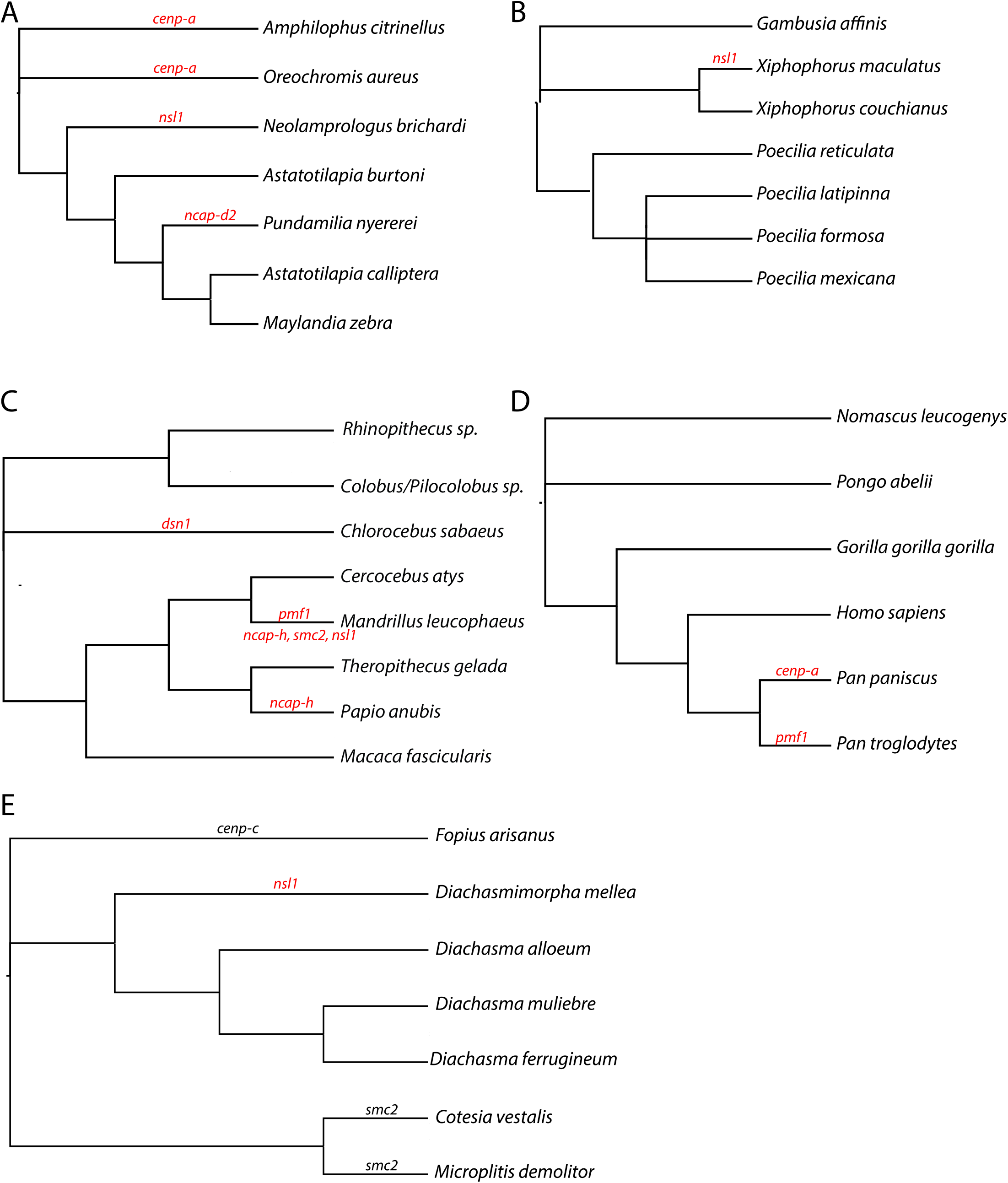
Phylogeny of each major animal group with missing genes in individual species indicated. Species trees of all species included in each animal group. Missing genes from each analysis are shown on individual branches. Red indicates they are confirmed missing from genome annotations. Black indicates they were removed for other filtering reasons.

### Multiple Alignments

Depending on the source of the sequence files, we obtained annotations either from Ensembl v98, NCBI, or from those published with original genome sequences (Tvedte et al. 2019; Tvedte et al. 2020) and trimmed sequences to include only the coding sequence (CDS). Amino acid sequence lengths are reported as the shortest sequence as missing sites are removed from all subsequent analyses (Summarized in Table S2). We performed multiple alignments using default parameters via the MAFFT v7.308 in Geneious v9.1.8 (www.geneious.com), the MUSCLE translation align function in Geneious build 2023-04-03 (www.geneious.com), and Seaview 5 for insects, fish, and primates respectively. Alignments were inspected and edited by eye in Geneious to remove gaps and stop codons.

### Tests of Positive Selection

To identify evidence of positive selection we used two groups of approaches within PAML suite version 4.9 (Yang 2007). First, we used the site-model test in the codeml program, using a tree-based approach to fit the data to the nearly neutral models (M7: Nsites =7) and a model that incorporates positive selection with a beta distribution with a fraction of sites ω > 1 (M8 Nsites =8) both with ω = 0.2. The fit of the models was compared using a log-likelihood ratio test (df =2). To assess if any individual sites were evolving under positive selection we used the Bayes Empirical Bayes (BEB) calculation of posterior probabilities for site classes (Yang 2005). Second, to generate hypotheses we estimated dN/dS (ω) for each branch independently using the free ratio model (Model = 1, Nsites = 0, fix_omega = 0, omega = 0.2) using a cutoff of ω > 1 as suggestive of positive selection. The free ratio model is very parameter rich and is not ideal for testing positive selection. We therefore expanded this analysis to branches exhibiting ω > 1 using the branch site model. The branch site model (Model = 2, Nsites = 2) allows omega to vary among sites and across branches and assumes that the phylogeny is divided *a* priori into foreground and background lineages. Using this test, we compared the neutral null model (fix_omega = 1 and omega = 1) to the alternative model of positive selection (fix_omega = 0 and omega = 1.5) on individual branches.

### Filtering false positives

There are several reasons an ω value may be elevated above 1 and not indicate positive selection in the free ratio model. We employed additional screening to remove these genes from our list of those evolving adaptively. First, individuals may exhibit an ω > 1 because of few or no changes. If dS = 0, this can inflate the dN/dS ratio which may result in a false positive. For example, if there is only a single nonsynonymous change, and dS = 0, then ω = 999 but this result is incongruous with positive selection and should be called a false positive. If, however, dS = 0 but there are substantial nonsynonymous changes in the protein, the estimated value of ω will be unreliable (ω = 999) but still consistent with positive selection. Another scenario resulting in a false positive is when only a single synonymous and single nonsynonymous change are present (e.g. dN = 0.000002, dS = 0.000001, dN/dS=2). Again, although ω > 1, the data are not consistent with positive selection and should be called a false positive. False Positives are indicated in Table S3.

### Species Trees

Species relationships were based on species trees available on Ensembl (accessed 12/2019-08/2020) for Cichlidae and Poeciliidae, Cercopithecidae, and Hominidae, and on an ASTRAL (v5.7.3) species tree inferred from BUSCO single-copy orthologs for Braconidae species in this study (see also (Tvedte et al. 2017)). Species tree figures were generated using FigTree version 1.4.4 (http://tree.bio.ed.ac.uk/software/figtree/).

### Protein complex structures

Schematic figures were developed based on structures from the Protein Data Bank including the CENP-A/CENP-C/Nucleosome Complex PDB:4X23 (Kato et al. 2013), the MIS12/CENP-C Complex PDB: 5LSK (Petrovic et al. 2016), and the Condensin I Complex. As the Condensin I full structure has not been resolved, we show this in two separate structures (1) including all subunits except NCAP-G PDB: 6YVU (Lee et al. 2020) and (2) NCAP-G and NCAP-H partial structure PDB: 5OQQ (Kschonsak et al. 2017). Model rendering was performed using UCSF ChimeraX (Meng et al. 2023).

## Results and Discussion

Rapid evolution of centromeres is pervasive across the tree of life (Henikoff et al. 2001; Malik and Henikoff 2001; Bayes and Malik 2008; Brown and O’Neill 2014). Rapid evolution of centromere-associated inner kinetochore proteins is also pervasive (Cooper and Henikoff 2004; Talbert et al. 2004; Schueler et al. 2010; Fountain and Kral 2011; Abbey and Kral 2015; Finseth et al. 2015; Kral 2016; Kumon et al. 2021). However, the prevalence of positive selection in other associated protein complexes beyond the inner kinetochore - including outer kinetochore and condensin complexes - warrants further investigation.

We chose to investigate genes encoding two protein complexes - Condensin I and Mis12. We selected these complexes based on several criteria. First, these protein complexes have different associations with the centromere through different inner kinetochore proteins - CENP-A and CENP-C respectively. The difference in association to the centromere was deliberate as previous studies, in multiple plant and animal species, showed evidence of positive selection often in either *cenp-c* or *cenp-a* but not always both (Talbert and Henikoff 2022). We also know binding domains and recognition sequences of kinetochore proteins are different across taxa (Kyriacou and Heun 2023), so it is possible that compensatory evolution between centromere-protein or protein-protein domains would also differ across taxa. Specifically, differences in binding domains would lead to differences in compensatory coevolution and different cascades of selection through protein complexes. Second, we selected these complexes because they were of similar size – in terms of number of protein subunits. The Condensin I complex contains five subunits (NCAP-G, NCAP-H, NCAP-D2, SMC2, SMC4) and the Mis12 Complex contains four subunits (MIS12, DSN1, NSL1, PMF1) (Figure 1). We know genes encoding Condensin I proteins previously showed evidence of positive selection in *Drosophila* (Beck and Llopart 2015) and genes encoding Mis12 complex previously showed evidence of positive selection in primates (Pontremoli et al. 2021), however systematic assessment of these complexes across groups had not been performed. We hypothesized that signatures of positive selection would be present in a subset of these genes in all tested groups of Metazoan animals. We also included an asexual diploid the amazon molly which should not be experiencing centromere drive. The hypothesis is that we would not see elevated dN/dS or signatures of positive selection in kinetochore proteins in this species.

We first used the codeml program in the PAML suite to identify signatures of positive selection in each animal group. Animal group sizes ranged; in Braconidae, Cichlidae, and Poeciliidae we used a maximum of seven species, in Hominidae we used a maximum of six species, and in Cercopithecidae we used a maximum of eight species. Comparing the fit of the nearly neutral model (M7) to a model that incorporates positive selection (M8), we found a significantly better fit of the M8 selection model in three genes in *Braconidae* (*ncap-d2, ncap-h*, and *pmf1*), two in Poeciliidae (*cenp-a* and *smc2)*, one in Cichlidae (*ncap-d2*), two in Hominidae (*ncap-g* and *nsl1*), and one in Cercopithecidae (*mis12*) (Table 1; Table S4).

**Table 1.**
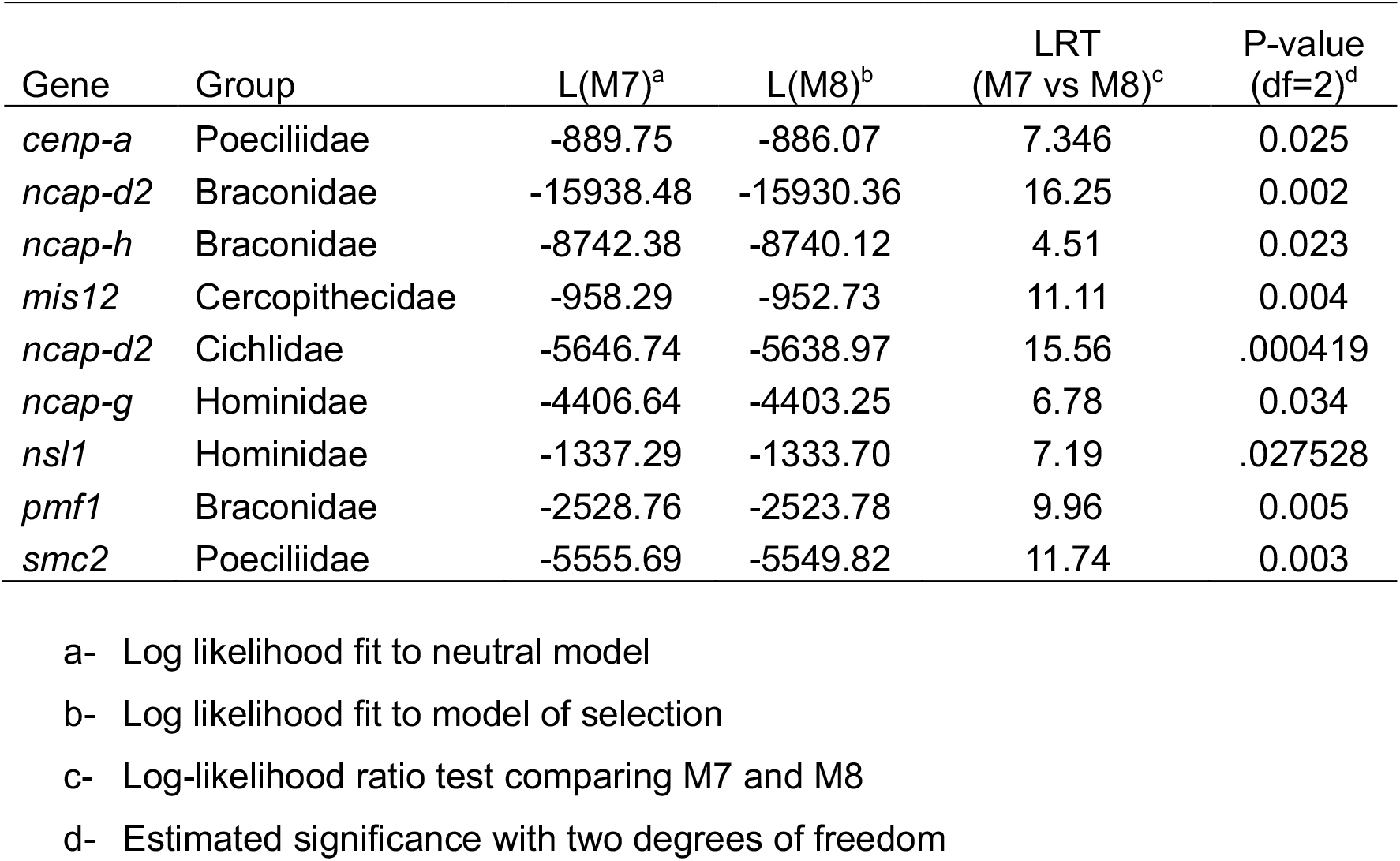
Site Model Test comparing M7 (Neutrality) to M8 (selection). Only significant results are shown.

To generate hypothesis, we calculated the number of nonsynonymous (dN) and synonymous (dS) changes per site in the coding region to identify genes with dN/dS (ω) > 1; indicative of potential positive selection for each branch (Table S3). After screening ω values for false positives (See Methods), we found widespread elevation of ω > 1. Specifically, in the inner kinetochore we found ω > 1 in *cenp-a* in both groups of fish and primates (we did not have reliable *cenp-a* sequences from Braconidae and did not include them in the analysis). These included: Cichlidae *(Pundamilia nyerei)*, Poeciliidae (*Gambusia* affinis, Xiphophorus *couchianus*, and *Poecilia mexicana*), Cercopithecidae *(Rhinopithecus bieti)*, and Hominidae (*Pongo abelii*) and in *cenp-c* in primates only (no *cenp-c* sequences could be found in either group of fish). These included: Cercopithecidae *(Rhinopithecus roxellana, Chlorocebus sabaeus, Cercocebus atys, Theropithecus gelada)* and Hominidae (*Gorilla gorilla gorilla* and *Pan paniscus*) (Figure 2; Table S3). These findings were consistent with previously published studies including evidence of positive selection in *cenp-a* in fish and evidence of positive selection in *cenp-a* and *cenp-c* in primates (Talbert and Henikoff 2022).

**Figure 2.**
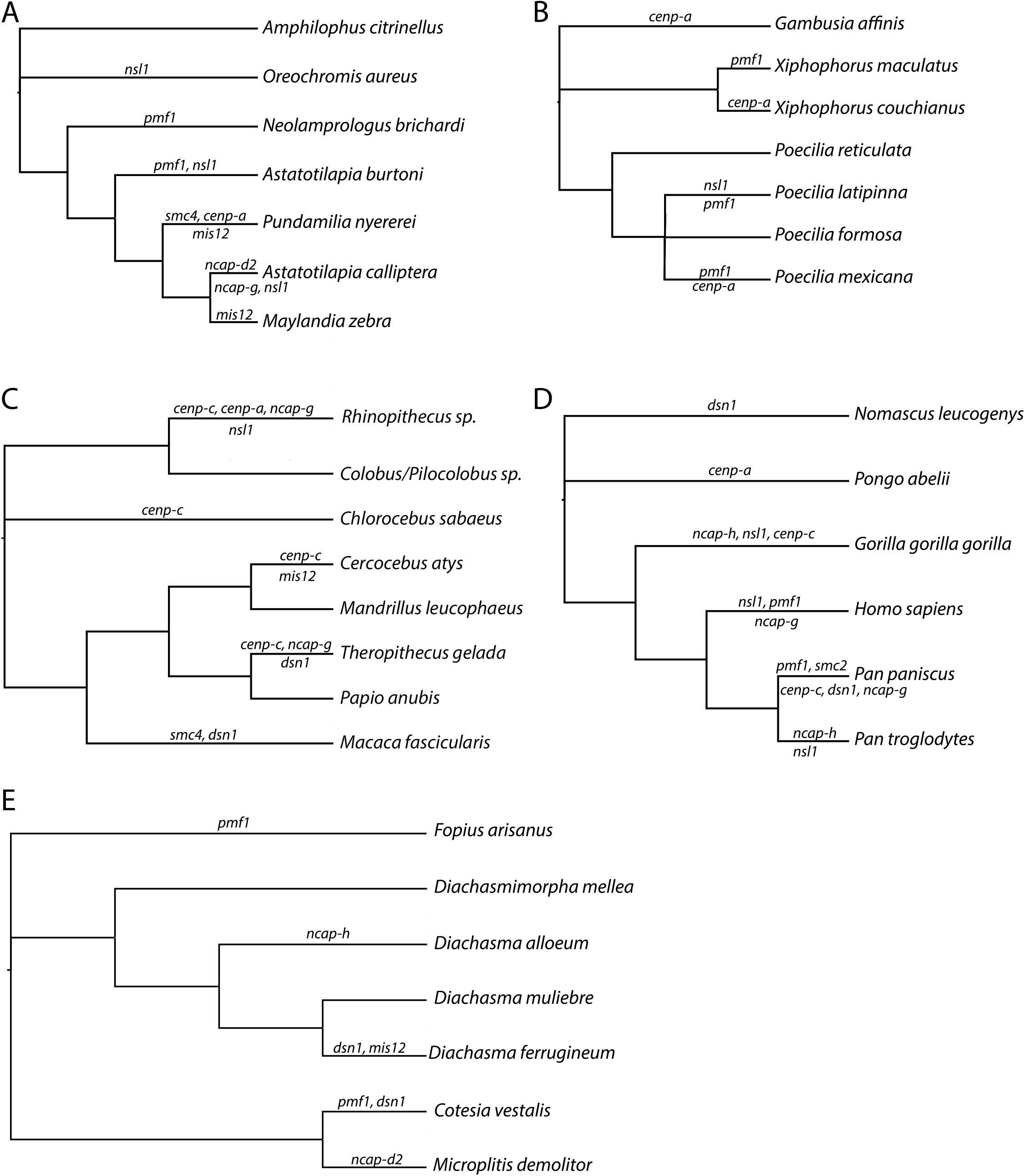
dN/dS branches labeled.
Species trees of all species included in each animal group with all genes exhibiting a dN/dS >1 indicated on each branch. These genes pass all filters described in the methods. (A) Cichlidae (B) Poeciliidae (C) Cercopithecidae (D) Hominidae (E) Braconidae

Beyond the inner kinetochore we also found many lineages with ω > 1 for one or multiple genes (Figure 2). In the Condensin I complex (SMC2, SMC4, NCAP-G, NCAP-H, NCAP-D2) we did not find ω > 1 in Poeciliidae. We did, however, identify ω > 1 in several genes encoding these proteins; Cichlidae: *ncap-g* and *ncap-d2* in *Astatotilapia calliptera* and *smc4* in *Pundamilia nyererei*; in Cercopithecidae: *ncap-g* in *Rhinopithecus bieti* and *Theropithecus gelada* and *smc4* in *Macaca fascicularis*; in Hominidae: *ncap-g* in *Homo sapiens* and *Pan paniscus* and *ncap-h* in *Gorilla gorilla gorilla* and *Pan troglodytes*; and in Braconidae: *ncap-h* in *Diachasma alloeum* and in *ncap-d2* in *Microplitis demolitor*. In the Mis12 Complex (MIS12, DSN1, PMF1, NSL1), we identified ω > 1 in several genes encoding these proteins; Cichlidae: *nsl1* in *Oreochromis aureus, Astatotilapia burtoni*, and *A. calliptera, pmf1* in *Neolamprologus birchardi* and *A. burtoni*, and *mis12* in *Pundamilia nyererei* and *Maylandia zebra;* in Poeciliidae *nsl1* in *Poecilia latipinna, pmf1* in *Xiphophorus maculatus, P. latipinnia*, and *P. mexicana;* in Cercopithecidae: *nsl1* in *Rhinopithecus roxellana, dsn1* in *Theropithecus gelada* and *Macaca fascicularis*, and *mis12* in *Cerocebus atys*; in Hominidae: *nsl1* in *Gorilla gorilla gorilla, Homo sapiens*, and *Pan troglodytes, dsn1* in *Nomascus leucogenys* and *P. paniscus*, and *pmf1* in *Homo sapiens* and *P. paniscus;* in Braconidae: *dsn1* in *Diachasma ferrugineum* and *Cotesia vestalis, pmf1* in *Fopius arisanus* and *C. vestalis*, and *mis12* in *D. ferrugineum* (Figure 2; Table S3). We did not identify ω > 1 in any gene in the meiotic diploid *P. formosa*.

Detecting ω >1 is good for generating hypotheses, but we also employed the branch site model tests to confirm positive selection. Negative results in branch site model tests do not mean there is no positive selection, instead the branch site model can confirm positive selection. In performing these confirmatory analyses, we lost substantial signal for positive selection (Table S5). In total, the branch site model only confirmed two cases of positive selection; *ncap-*d2 in *A. calliptera* and *cenp-a* in *R. bieti*.

Our results showed positive selection via the site model test and indicated potential selection detected by dN/dS (ω >1) in many genes across our selected groups of Metazoans. In the inner kinetochore, our results mirrored those of others, identifying positive selection in *cenp-a* in fish and in *cenp-a* and *cenp-c* in primates (Talbert and Henikoff 2022). However, in most cases, potential positive selection indicated by ω >1 did not confirm in follow-up branch site model tests and we did not identify widespread positive selection as was reported previously in *Drosophila* (Beck and Llopart 2015; King et al. 2019). However, we only had single sequences per species available for our analyses and without polymorphism data we are limited in our tools to identify signatures of positive selection. We specifically could not employ methods for identifying repeated long-term positive selection (McDonald and Kreitman 1991) or selective sweeps (Smith and Haigh 1974). This means we are likely underestimating the prevalence of positive selection across these gene groups.

Interestingly, we never found ω >1 in any gene in *P. formosa* the asexual diploid that we hypothesized would not experience centromere drive or compensatory positive selection in kinetochore proteins. Still, absence of evidence is not evidence of absence and more exploration into these dynamics in asexual diploids is warranted. One additional limitation in understanding how centromere to kinetochore dynamics impacts compensatory evolution, is the lack of knowledge about specific protein-protein interaction domains across taxa. We know inner kinetochore protein binding domains differ across species and at times the protein content is different across taxa. Therefore, more work in this area could help with analyses of centromere-kinetochore compensatory coevolution.

## Acknowledgements

We would like to thank our funding sources including the National Institutes of Health, National Institute of General Medical Sciences NRSA fellowship F32GM122419 to E.A.B and start-up funds from University of Kansas Research Rising to E.A.B.

## Figure and Supplemental Material Legends

***Supplemental Table 1***. Accession IDs for sequencing data.

***Supplemental Table 2***. Gene names and lengths.

Lengths are reported as the shortest sequence as missing sites are removed from analysis via PAML.

***Supplemental Table 3***. dN/dS of all branches estimated from the free ratio model.

***Supplemental Table 4***. Site Model Test comparing M7 (Neutrality) to M8 (selection). Includes significant and non-significant results.

***Supplemental Table 5***. Branch Site Model Test for all terminal branches with dN/dS > 1.

